# Terasaki Spiral Ramps and Intracellular Diffusion

**DOI:** 10.1101/675165

**Authors:** Greg Huber, Michael Wilkinson

## Abstract

The sheet-like endoplasmic reticulum (ER) of eukaryotic cells has been found to be riddled with spiral dislocations, known as ‘Terasaki ramps’, in the vicinity of which the doubled bilayer membranes which make up ER sheets can be approximately modeled by helicoids. Here we analyze diffusion on a surface with locally helicoidal topological dislocations, and use the results to argue that the Terasaki ramps facilitate a highly efficient transport of water-soluble molecules within the lumen of the endoplasmic reticulum.

## 1. Introduction

The endoplasmic-reticulum (ER) sheets consist of stacks of pairs of phospolipid bilayer membranes in the interior of eukaryotic cells [1]. The bilayers divide the cell into two distinct regions, illustrated in figure 1(a): the lumen of the ER enclosed between the doubled bilayers forms a single region which is connected throughout the cell, and which is also continuous with the nuclear envelope. The region complementary to the lumen is the cytoplasm (and also the nucleoplasm). As well as dividing the cell into two regions, the surfaces of the ER play an important role in organizing complex biochemical processes by acting as a substrate for membrane-bound protein complexes on both the luminal and cytoplasmic sides (*e.g.*, ribosomes are attached to the cytoplasmic facing ER-sheet membranes). It is this role as a surface for catalysis that dictates the large surface area of the ER, but these extensive surfaces would create barriers to diffusion in both compartments [2]. Specifically, a system of stacked bilayer membranes can act as a barrier to diffusion of water-soluble species in the directions perpendicular to the layers. This appears to present a challenge to the efficient operation of the cell.

**Figure 1.**
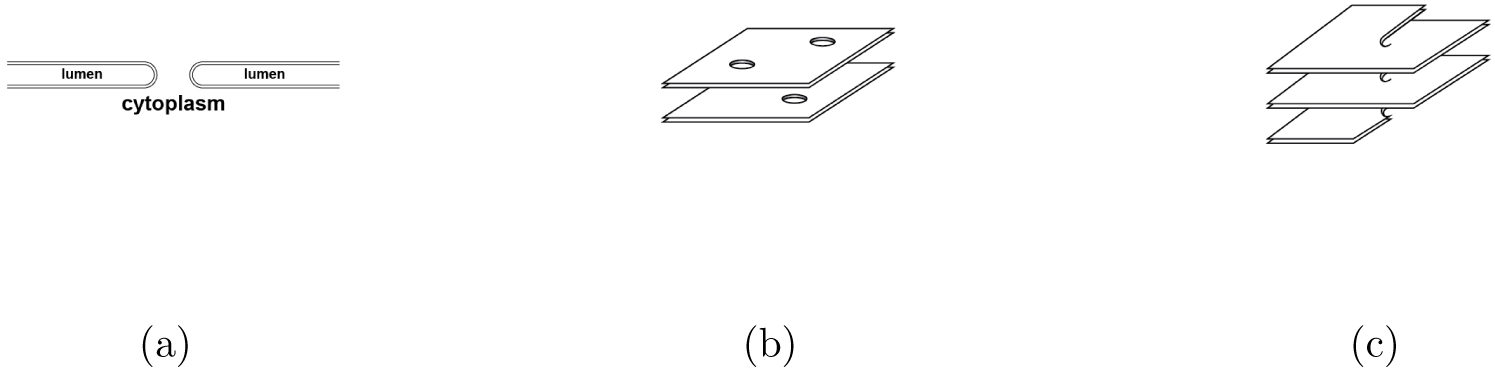
The interiors of eukaryotic cells are divided by the surfaces of the endoplasmic reticulum (ER). (**a**) Each of these surfaces contains a region termed the ‘lumen’, separated from the cytoplasmn by lipid bilayers. Edges of the ER may have a catenoidal cross-section. (**b**) The ER is sometimes pictured as being punctured by ‘windows’, allowing diffusion of water-soluble species throughout the cytoplasm. (**c**) However, careful analysis of serial-section transmission-electron micrographs reveals the presence of spiral dislocations termed *Terasaki ramps*.

It has been understood for some time that there are topological ‘defects’ in the layers of the ER. These are often represented as holes in the lipid bilayer system, with approximately catenoidal edge surfaces, as illustrated in figure 1(a), forming a set of ‘windows’ or fenestrae, as illustrated in figure 1(b). However, careful studies [3, 4] of the topological structure of the ER sheets have revealed that the layers have a type of screw dislocation, which have been named ‘Tersaki spiral ramps’ after their primary discoverer. These are illustrated schematically in figure 1(c). In this paper we argue that these spiral dislocations allow extremely efficient diffusive transport perpendicular to the plane of the lipid bilayers.

Small molecules are able to traverse the cell by diffusion, a consequence of the fact that the motion of each molecule is a random walk. For both fenestra and Terasaki ramps, a random-walk trajectory can allow a molecule to pass between regions of the cytoplasm separated by sheets of the ER. In the case of a window connecting layers of the ER (as illustrated in figure 1(b)), a molecule must diffuse to the aperture in order to pass between layers (as illustrated in figure 2(a)). In the case of a spiral dislocation, however, a path (such as that illustrated in figure 2(b)) which winds once around a spiral dislocation allows movement between two successive sheets of the endoplasmic reticulum, without having to make contact with the dislocation itself. Compared to the situation illustrated in figure 2(a), this represents a much weaker constraint on the subset of random walks which allow transport between sheets. For this reason, we argue that, rather than being a structural curiosity, the Terasaki spirals may be an essential element in the efficient operation of eukaryotic cells.

**Figure 2.**
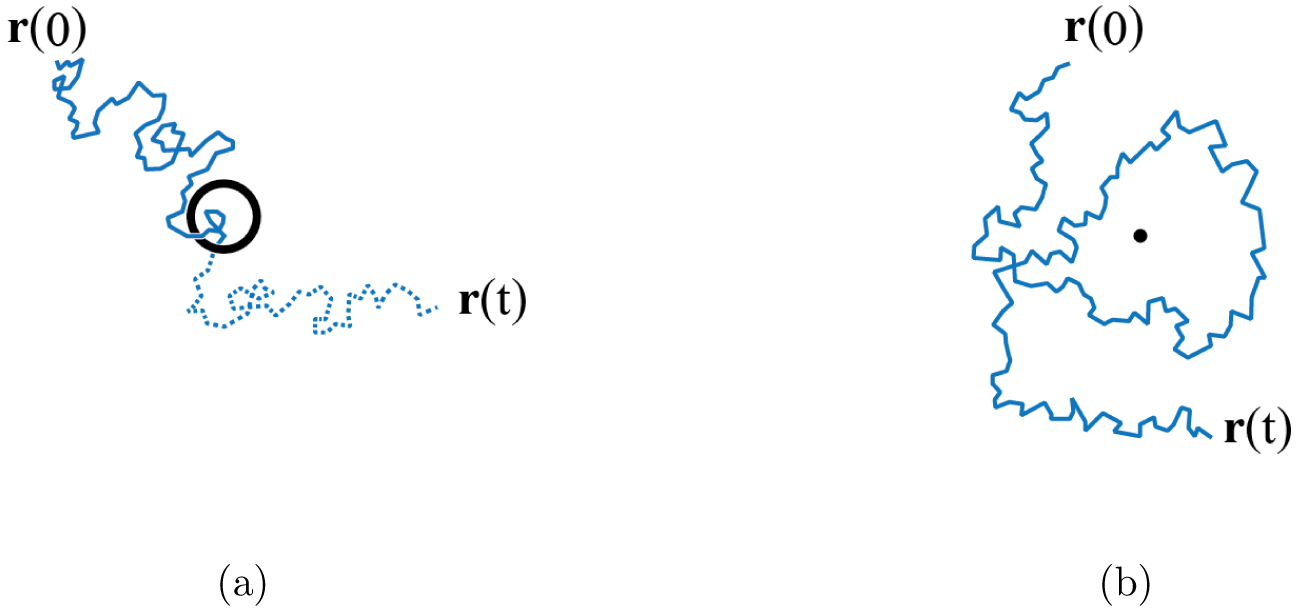
Diffusion is a consequence of molecules following a random walk trajectory. Some of these trajectories allow crossing of the ER sheets. In the case of a ‘window’, the trajectory has to make contact with the singularity (**a**), whereas in the case of a spiral dislocation, the trajectory only needs to wind around the singularity (**b**).

The technical content of our paper is concerned with describing a model for diffusion in the endoplasmic reticulum, showing how the spiral dislocations allow efficient transport perpendicular to its layers. In section 2 we model the structure of the endoplasmic reticulum in the vicinity of a spiral dislocation as a helicoidal surface. We analyze the diffusion on a helicoidal surface, emphasizing the statistics of winding numbers of diffusive trajectories about its axis. In section 3 we show that, at large times, the statistics of winding numbers approach those of random walks in a plane, which avoid a disc centered on the position of the dislocation. We calculate the variance of the winding number at large times. Having modeled a single dislocation, in section 4 we use our results to describe diffusion of a water-soluble molecule in the cell, modeling the endoplasmic reticulum as a set of sheets connected by spiral dislocations with an approximately helicoidal structure. Section 5 discusses the implications of our estimate. The dispersion is described by an effective diffusion coefficient which is proportional to the density of Terasaki ramps. We argue that the density of ramps that are present in the ER is sufficient to allow unimpeded diffusion of small molecules.

Our results are related to a previous treatment of diffusion in an extended lamellar medium with randomly scattered dislocations [5]. That earlier work predicted that dispersion perpendicular to the plane of the lamellae is marginally super-diffusive (specifically, the variance increases faster than linear in time by a logarithmic factor). However, the prediction of super-diffusion was strongly criticized [6], and the issue was left unresolved. In section 5, and in an Appendix, we explain why the criticisms are not relevant to our application.

## 2. Diffusion on a helicoid

The Teraski spiral ramps are screw dislocations connecting the layers of the endoplasmic reticulum. The axis of the spiral is a topological singularity, in the sense that if a path on the surface of the endoplasmic reticulum lipid bilayer makes a circuit about the axis, then the path ends up on another layer of the structure. The surface of the lipid bilayer structure is, however, smooth everywhere. A helicoid is a smooth surface which has the same topology as the Terasaki spiral. The Helfrich model [7] for the energy density of a biological membrane has a term proportional to the square of the mean curvature. Because the helicoid has zero mean curvature it is a plausible model for the shape of the Terasaki spirals: a more refined model is considered in [8]. We therefore begin our investigation of diffusion within the endoplasmic reticulum by analyzing diffusion on a helicoidal surface.

A helicoid is a two-dimensional surface in three dimensions defined by the following parametric equations for the Cartesian coordinates (*x, y, z*):

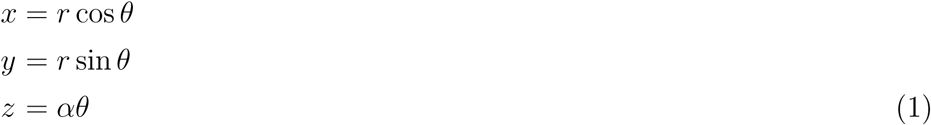

with *r* ≥ 0 (technically speaking, it is the *half helicoid* since the line *r* = 0 represents an edge of the structure). We assume there is isotropic diffusion on the two-dimensional surface, with diffusion coefficient *D*. There is a corresponding stochastic dynamics of the *r, θ* variables. Because our particular interest is in the dynamics of the *z* coordinate, representing motion perpendicular to the sheets, we concentrate on analyzing the dynamics of *θ* in the limit as time *t* → ∞. For simplicity, in this section we consider diffusion in a cylinder of radius *R*, with the dislocation lying along the axis of the cylinder (equivalently, we consider the motion to occur in the region 0 ≤ *r* ≤ *R*, with *θ* unbounded.)

The problem of diffusion on a helicoidal surface is closely related to understanding the winding of a random walk about a point in the plane. Some classic works which address the distribution of winding numbers for diffusion on the plane are [9, 10, 11, 12, 13]. The plane-diffusion problem turns out, however, to be quite different, in that the winding number about a point has a long-tailed distribution with a diverging variance. For our problem of helicoidal diffusion, we show that the winding-number variance is finite.

We can introduce a local Cartesian frame describing points on the helicoid in the neighborhood of (*r, θ*), with coordinates (*X, Y*). Because of rotational symmetry it is sufficient to take *θ* = 0. The *X* axis will be taken to lie in the radial direction, along a line of constant *z*, and the *Y* axis is then tilted so that small increments *δy, δY* of *y* and *Y* are related by 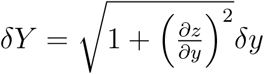. Noting that d*z* = *α*d*θ* and d*y* = *r*d*θ*, we have

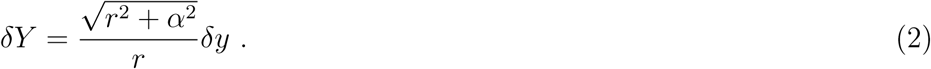

Because a point makes diffusive motion on the helicoidal surface, we can assume that, in a small time *δt*, there is a corresponding diffusive motion on the tangent plane. The consequent small displacement on the tangent plane, (*δX, δY*), has a probability density function (PDF) which is the diffusion kernel for isotropic diffusion in two dimensions:

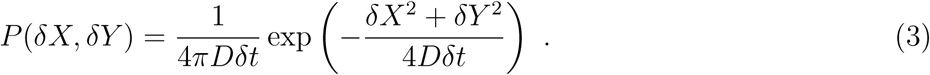

This implies corresponding random displacements of the parameters *δr* and *δθ*. If we can determine the first two moments of these displacements, we can write down a Fokker-Planck equation for the joint PDF *𝒫*(*r, θ, t*) of *r* and *θ*:

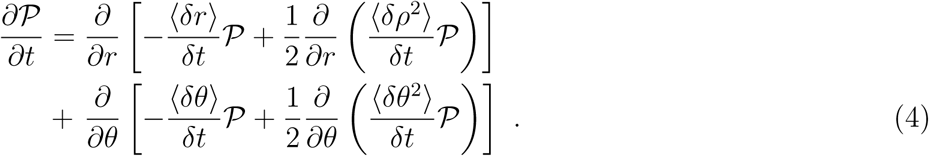

The relations between (*X, Y*) and (*r, θ*) are determined by first projecting onto the (*x, y*) plane, then transforming to polar coordinates. We have *δx* = *δX* and 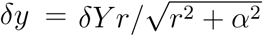. Then, noting that *r*^2^ = *x*^2^+*y*^2^ and using 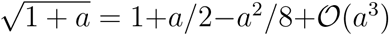, retaining terms to second order in the stochastic fluctuations, we have

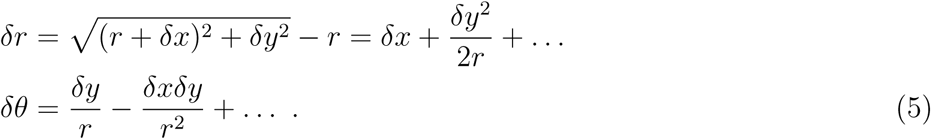

Hence we find

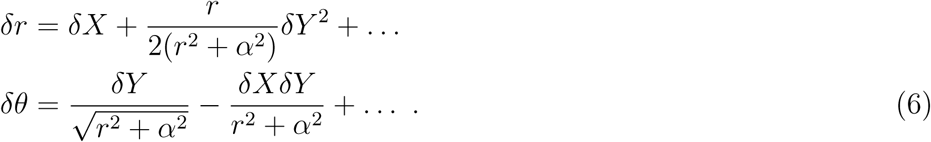

Noting that ⟨*δX*^2^⟩= ⟨*δY*^2^⟩= 2*Dδt* and ⟨*δX*⟩ = ⟨*δY* ⟩ = ⟨*δXδY* ⟩ = 0, the statistics of the increments of the polar coordinates are, therefore,

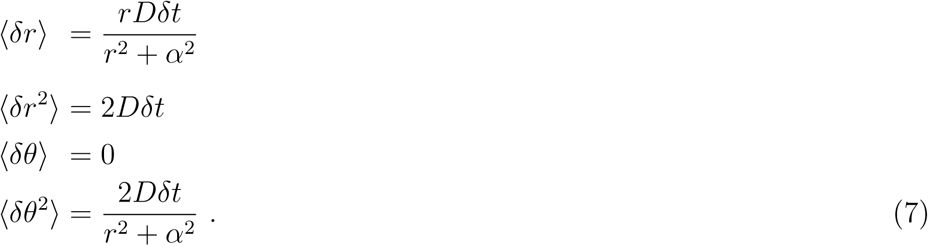

The Fokker-Planck equation is therefore

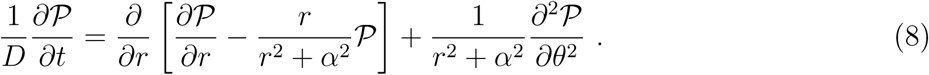

Now consider the long-time behavior of the distribution of the angular variable, for the case where the motion is confined to a cylinder with radius *R* with the axis of the helicoid at its center. We assume that the radial distribution has reached equilibrium, and write

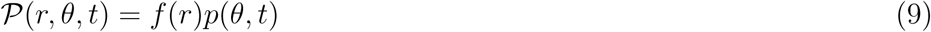

where the radial component is the zero-flux steady-state solution of (8), satisfying *f* ′ = *fr/*(*α*^2^ + *r*^2^) with solution

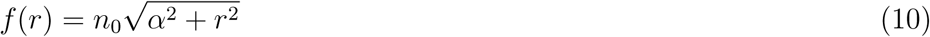

where *n*_0_ is a constant of integration. Noting that a uniform distribution on the disc of density *n*_0_ corresponds to *f* (*r*) = *n*_0_*r*, we see that *n*_0_ can be identified with the density when *r/α* ≫ 1. We assume the distribution *p*(*θ, t*) is normalized, so that its integral over *θ* is equal to unity. If there is a single particle in a disc of radius *R*, then 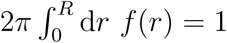, so that when *R/α* ≫ 1, the density is *n*_0_ ∼ 1*/πR*^2^.

Now consider the long-time behavior of *θ*. The statistics of the increments of *θ* are specified in equations (7). Because the variance of *δθ* depends upon *r*, the long-time behavior of ⟨*θ*^2^⟩ is determined by averaging over the distribution of *r*. The distribution of *θ* has, in the long-time limit, a variance 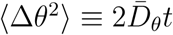, with diffusion coefficient

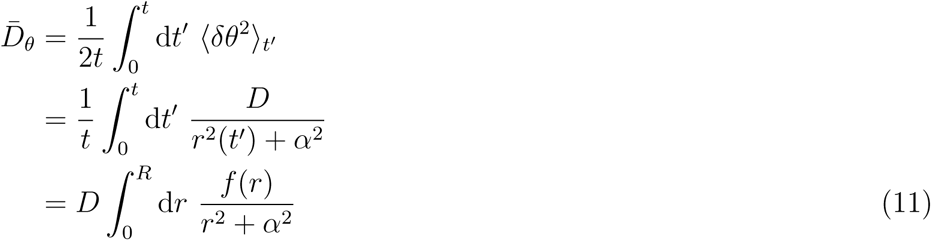

where *R* is the radius of the cylinder. The last step of (11) follows from replacing a time average with an ensemble average, and using the fact that *f* (*r*) is the probability density function for *r*. Hence we find

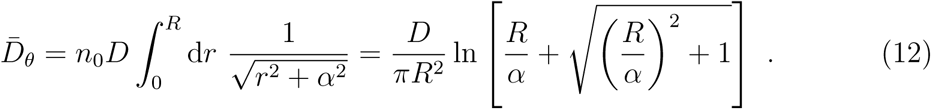

In the case where *R/α* ≫ 1, this is

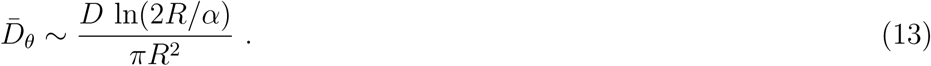

## 3. Winding number at a finite time

In section 2 we examined the diffusion of the rotation angle in a finite region, and used the idea that the diffusing particle approaches a uniform density at large times. In this section we consider how to evaluate the variance of the rotation angle in an unbounded region.

When *r* ≫ *α*, the Fokker-Planck equation (8) takes the same form as the two-dimensional diffusion equation. Because the diffusing particle has a low probability of being in the vicinity of the dislocation, this indicates that, for large time, the variance of the rotation angle will be determined by solving a conventional diffusion equation, with the center of the helicoid replaced by an impenetrable disc, with a radius *ϵ* which will be determined shortly.

Compare equation (13) with the case of diffusion in the plane around a disc of radius *ϵ*. The corresponding angular diffusion coefficient is obtained by setting *α* = 0, and introducing a lower cutoff of *ϵ* in the integration over *r* in (11). This gives an equation which is identical to (13), except for replacing *α/*2 with *ϵ*, implying that a helicoid with pitch *α* has the same asymptotic winding number statistics as a planar diffusion around a disc of radius *α/*2.

We have argued that the diffusion on a helicoid of pitch *α* is equivalent to motion on a flat surface with an excluded disc of radius *ϵ* = *α/*2 for trajectories which do not approach close to the axis of the helicoid. Now we use this observation to determine the distribution of winding angle Δ*θ* when the starting point is at a distance *R* from the axis, with *R/α* ≫ 1.

The winding angle of a trajectory is

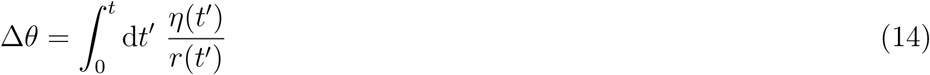

where *r*(*t*) is the distance of the diffusing trajectory from the center of the disc at time *t*, and *η*(*t*) is a stochastic velocity of the diffusive trajectory in a direction perpendicular to its displacement from the dislocation. This satisfies

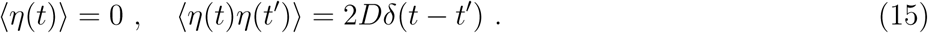

Using (15) in (14), the variance of the rotation angle is

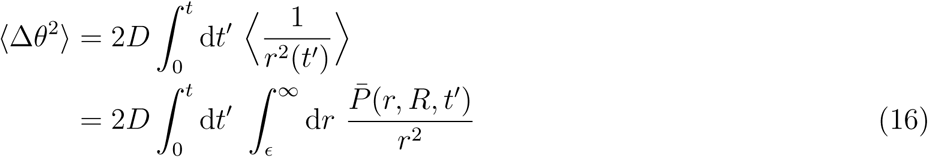

where 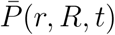 is the probability density to reach a distance *r* from the dislocation at time *t*, after starting at *R* when *t* = 0. Integrating the propagator for diffusion in two dimensions (equation (3)) over a circle, we obtain

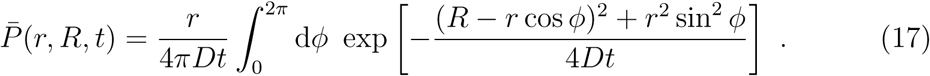

The variance of the rotation angle at time *t* for a trajectory which starts at a distance *R* from the dislocation is therefore

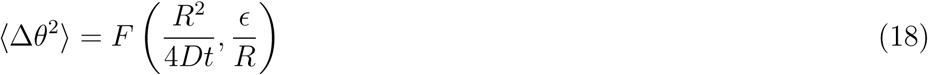

where

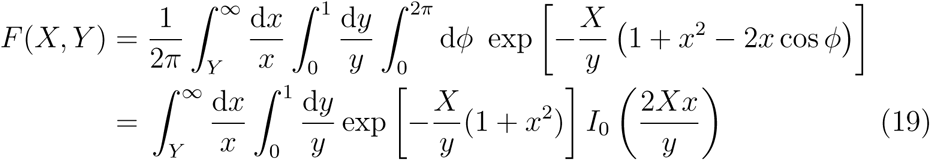

and *I*_0_(·) is a modified Bessel function of the first kind and of order zero. This may be written in the form

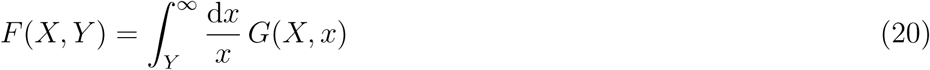

where *G*(*X, x*) is obtained by comparison with (19).

Let assume that *Y* ≪ 1, and divide the integral over *x* into two intervals: [*x*_0_, 1] and [*Y, x*_0_], with *Y < x*_0_ ≪ 1. If *G*(*X*, 0) ≠ 0, in the limit as *Y* → 0, the integral in (19) is dominated by the second of these contributions, and we may write

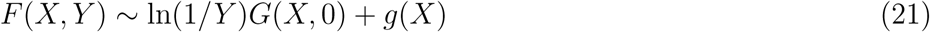

where *g*(*X*) can be expressed as a limit of a triple integral. The dominant term, logarithmic in *Y*, is proportional to

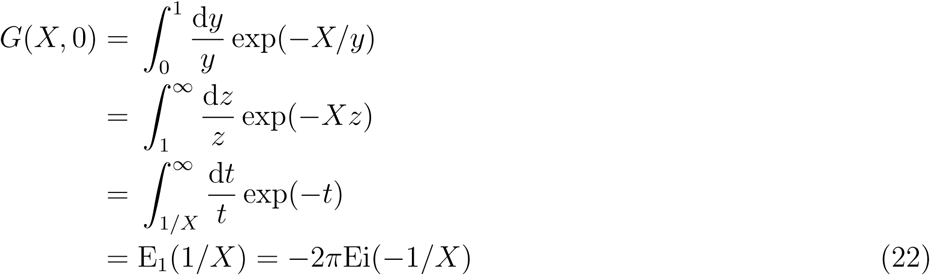

where *E*_1_(*z*) and Ei(*z*) are different standard specifications of the exponential integral function.

Recalling that *ϵ* = *α/*2, in the limit where *R* ≫ *α* the variance of the winding number is therefore approximated by

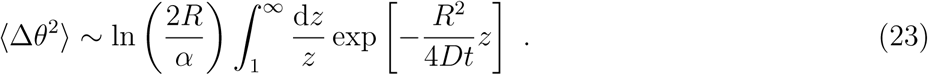

## 4. Model for perpendicular diffusion

We have analyzed diffusion on a helicoidal surface, leading to an estimate (23) for the winding angle of a trajectory after time *t*, starting at a distance *R* from the dislocation. We now adapt the results to model diffusion perpendicular to the sheets of the endoplasmic reticulum, using the observation that its sheets are connected by multiple spiral dislocations. For definiteness, we consider diffusion within the lumen of the ER, but the basic arguments for modeling diffusion in the cytoplasm differ only in inessential points. This perpendicular diffusion process is described by keeping track of on which sheet of the lumen a molecule is located. Because the spiral dislocation singularities connect different sheets to form a single manifold, its subdivision into numbered sheets is somewhat arbitrary.

The layers of the endoplasmic reticulum, which we assume have mean separation *h*, are connected by many of these dislocations, which will be assumed to be randomly scattered, with planar density *ρ*. The dislocations may be either ascending or descending for a positive winding number, and we distinguish these cases by a ‘charge’ *σ*_*i*_ = ±1 for the dislocation with index *i*. We shall assume that the distribution of these topological charges can be modeled by independently assigning each singularity a positive or negative charge, each with probability equal to one half. This net neutrality of topological charge is consistent with the census of Terasaki ramps from topological reconstruction of electron microscopy data [4].

We can make a specific model for the connected surface of the endoplasmic reticulum by considering the following function:

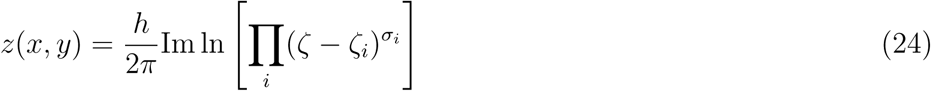

where *ζ* = *x* + i*y*, and where *ζ*_*i*_ are points randomly scattered in the complex plane, with the charges *σ*_*i*_ being randomly assigned to ±1 with probability 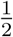. The spacing of the layers, *h*, is assumed to be small compared to the typical distance between nearest-neighbor dislocations. In the context of modeling the ER, this model has the attractive feature that the height *z* is a harmonic function. This implies that, in regions where the gradient of *z*(*x, y*) is small, the surface approximates a minimal surface (that is, the mean curvature is everywhere zero).

The motion of a molecule is now described by a random walk in the (*x, y*) plane, and its motion in the perpendicular coordinate (*z*) is determined by the manner in which this path is threaded through the set of singularities. If we were dealing with closed paths, we could describe the vertical motion by determining the winding number of the path about each singularity, but the diffusive trajectory of a molecule is almost always described by an open path. We adopt the following convention: From each dislocation we take a line parallel to the *x*-axis, in the negative direction. We regard the transitions between layers as occurring when a path crosses one of these lines. A path with decreasing *y*-coordinate moves up a layer as it crosses a line attached to a positive dislocation, and down a layer if it crosses a line attached to a negative dislocation (see figure 3). In the case where the path is closed, this convention is equivalent to summing the winding numbers of the trajectory about each dislocation, weighted by their sign.

**Figure 3.**
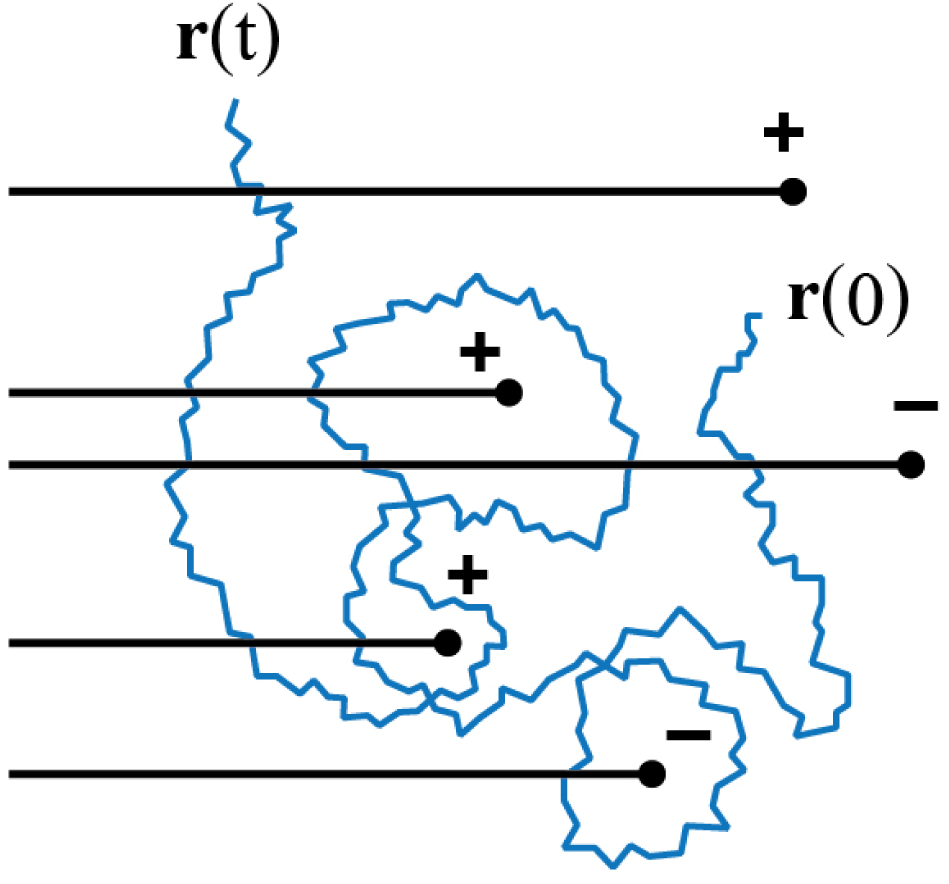
Our model of the endoplasmic reticulum, summarised by equation (24), has dislocations with random sign *σ*_*i*_ and random positions ***r****_i_*. A Brownian trajectory can move between ‘levels’ of the multi-sheeted manifold. We define a convention for labelling the levels of the manifold by the considering when the trajectory crosses a set of lines.

For a given path, we can define its winding number, *n*_*i*_, about a given dislocation with index *i*, in terms of the number of times the line attached to the singularity is crossed counterclockwise, minus the number of clockwise crossings. The change in level of a given path is the sum of the winding numbers for each dislocation, weighted by the sign of the dislocation:

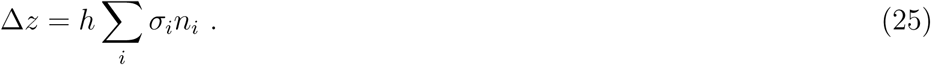

Note that the spacing between levels is *h* = 2*πα* for the simple helicoidal model studied in section 2. When time *t* is large, there will be many singularities which could have non-zero winding number, and the winding numbers will typically be large, so that we may approximate *n*_*i*_ = Δ*θ*_*i*_/2*π*, where Δ*θ*_*i*_ is the winding angle of the trajectory about the singularity with index *i*.

Both the winding numbers *n*_*i*_ and the signs of the dislocations *σ*_*i*_ are random variables (with zero mean), so that Δ*z* is a random variable. The distribution of the vertical displacement, Δ*z*, is conveniently described by its variance: for a fixed configuration of the signs *σ*_*i*_, this is

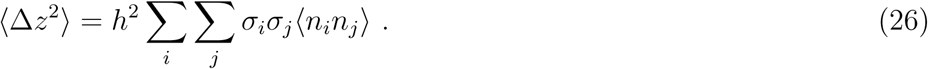

The correlation function of the winding numbers, ⟨*n*_*i*_*n*_*j*_⟩, can be computed, as discussed by Hannay [14]. However we shall perform a further average of (26) over the random signs *σ*_*i*_. Because ⟨*σ*_*i*_*σ*_*j*_⟩ = *δ*_*ij*_, the off-diagonal contributions to the double-sum are zero, so that the winding-number correlation function ⟨*n*_*i*_*n*_*j*_⟩ is not required for our calculation of ⟨Δ*z*^2^⟩. When we average over the signs, the doubly-averaged second moment of Δ*z* is

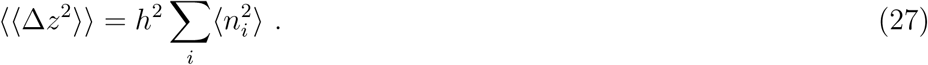

We have obtained the variance of the winding angle for a single helicoidal dislocation in equation (23). We now estimate the sum in (27) by replacing the sum by an integral over the density of dislocations at a distance *R* from a randomly chosen point. The expected number of dislocations in an annulus of width *δR* at distance *R* from the origin is

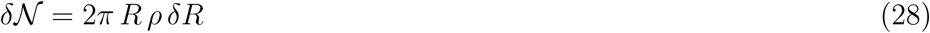

where *ρ* is the density of dislocations. Using equations (27), (23), (28) and recalling that *h* = 2*πα*, we have

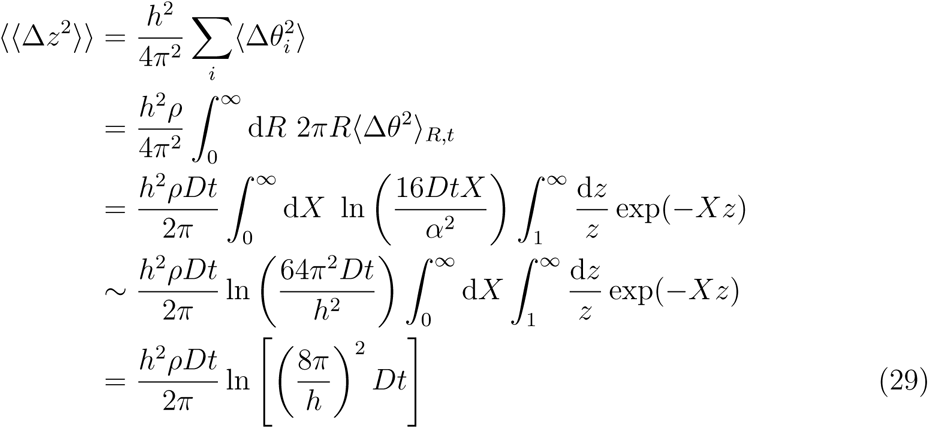

where in the penultimate step we use the fact that *Dt/h*^2^ ≫ 1 for times large enough to allow non-zero winding numbers with a significant probability. This is the principal technical result of our paper. A similar, but not identical, result has been proposed for diffusion in an extended lamellar medium punctured by dislocations [5]. We comment on the relation between these results in the concluding section. Note that ⟪Δ*z*^2^⟫ has a faster than linear growth, because of the logarithmic factor, implying that the dispersion in the *z* direction is marginally super-diffusive.

We can describe the dispersion across the layers of the ER by an effective diffusion coefficient, *D*_eff_. We define this by assuming that the size of the cell is 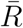, and noting that the time 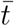 for dispersion by conventional diffusion may be related to 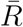 by writing 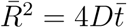. The effective diffusion constant for dispersion across the layers of the ER is defined by writing

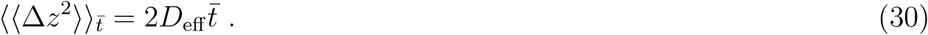

Using these definitions we find the effective diffusion coefficient perpendicular to the layers of the ER to be

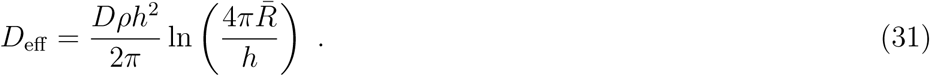

## 5. Discussion

We have argued that, relative to spiral dislocations, holes have a disadvantage when it comes to allowing perpendicular diffusive transport in the lumen and cytoplasmic space. This is because a dislocation allows perpendicular transport just by winding around the singular axis, whereas holes require that the trajectory has to go to the defect and pass through. This indicates that spirals allow very efficient perpendicular transport.

Our estimate for the effective trans-layer diffusion constant of our model, equation (31), differs from the aqueous diffusion coefficient *D* by a factor proportional to *ρh*^2^. Our model assumes that the singularities are distinct objects, which is equivalent to specifying that *ρh*^2^ ≪ 1, in which case the perpendicular diffusion coefficient is smaller than the planar diffusion coefficient.

Values for the inter-sheet separation in the ER are typically *h* ≈ 200nm. The spacing of of dislocations is thought to be roughly 1 *µ*m, hence we estimate that the small parameter is *ρh*^2^ ≈ 4 *×* 10^−2^. The characteristic size of the region occupied by the endoplasmic reticulum is comparable to the size of a cell, so 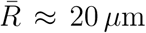 These estimates give

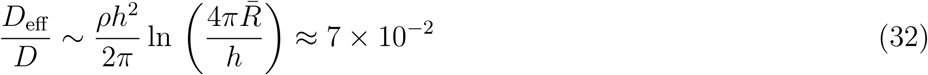

so that, while the perpendicular diffusion coefficient *D*_eff_ is smaller than the aqueous coefficient *D*, by a factor of approximately 15, it is still quite adequate to allow efficient transport of small molecules around the cell. If *D* = 10^−10^ m^2^s^−1^, which is typical for a small protein such as green fluorescent protein (GFP), the timescale to transverse the cell would be approximately ten seconds, which is negligible. Furthermore, measurements of diffusion of GFP in the endoplasmic reticulum [15] indicate that the diffusion coefficient is approximately ten to twenty times smaller than the diffusion coefficient in water, which is consistent with equation (32). We conclude that Terasaki ramps greatly facilitate transport of aqueous solutes through the eukaryotic cell. This naturally suggests the hypothesis that such topological ER structures are required for efficient diffusion in eukaryotes.

It is interesting to remark that, according to equation (29), ⟪Δ*z*^2^⟫ increases faster than linearly as a function of time, despite the fact that the underlying mechanism is diffusion on a complex surface. If we were modeling diffusion in an extended medium, rather than a finite-sized cell, this would pose a problem, because the predicted dispersion would eventually exceed that of the underlying diffusion process, which is impossible. (This point was made in a comment on the work by Gurarie and Lobkovsky [5], which treated diffusion in an extended lamellar phase [6, 16].) While this issue is not directly relevant to our estimates of diffusion in a cell, it is instructive to understand the origin of the difficulty, and how it could be resolved if we were dealing with an extended system. We address this in an appendix.

While our results show that small molecules can access the entire cytoplasm by diffusion alone, this may only be part of the ER-transport story. There is recent evidence for active transport throughout the smooth ER involving fluid flow within ER tubules. This may be required for the transport of larger proteins through the ER lumen, and hence to the most distal locations in cells [17].

## Acknowledgements

MW thanks the Chan Zuckerberg Biohub for its hospitality.

## 6. Appendix

If we are only interested in diffusion within a the dimensions of a typical cell, then the logarithmic term in equation (29) is of no concern, because *t* must become very large before the product of the small parameter *ρh*^2^ and the logarithmic factor exceeds unity. If we were interested in diffusion in a homogeneous region however, it is necessary to consider how the formulation of the problem should be modified.

Our model for the height of the surface, equation (24), is not suitable for describing an infinite, homogeneous region. To make the model well defined we have to confine the singularities *ξ*_*i*_ to a finite region, for example a disc of radius *ℛ*. The fluctuations of the product Π*_i_*(*ξ* − *ξ*_*i*_)^σ*_i_*^ increase without bound as *ℛ* increases. If we were interested in modeling an infinite region, we could replace (24) by

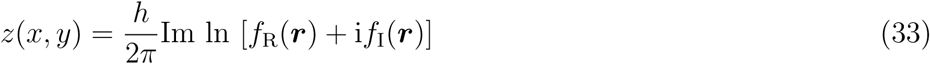

where ***r*** = (*x, y*) and *f*_R_ and *f*_I_ are independent realiizations of an ensemble of random functions. These functions can be assumed to have the following statistics:

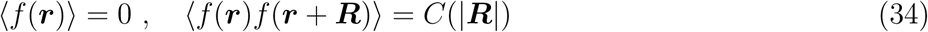

where *C*(*R*) is the correlation function of the random fields. This model is, by construction, statistically homogeneous, and is therefore suitable as a model for an infinitely extended random surface. (However it does lack the property of being a harmonic function, which was desirable for modeling the ER.) The dislocations correspond to points where *f*_R_ = *f*_I_ = 0. The density of these points is readily determined by the Kac-Rice method [18, 19].

At first sight, it seems as if the calculation in section 4 would be directly applicable to this variant model (equations (33) and (34). There is, however, a reason why the analysis does not carry over. In section 4, when we averaged over the distribution of signs *σ*_*i*_, we assumed that they are completely random. It has been shown that the distribution of zeros of a statistically homogeneous random field must satisfy a ‘screening’ property, implying that the signs cannot be chosen at random [20]. For this reason, the calculation of section 4 cannot be applied to the case of an infinitely extended region.

